# G-SPRI: A Structure-Centric Graph Model for Comprehensive Prediction of Cancer Driver Events from Missense Mutations

**DOI:** 10.64898/2026.05.06.723398

**Authors:** Boshen Wang, Ali M. Farhat, Bowei Ye, Jie Liang, Lei Yu, Zeyu Lu, Xinlei Wang, Lin Xu

**Author notes:** Corresponding Authors: Lin Xu, PhD.

## Abstract

*In silico* approaches for predicting the functional impact of missense mutations are critical for interpreting personal genomes and identifying disease-related biomarkers. Existing methods largely rely on sequence-based information or intuitive structural features, but often overlook the complex biophysical patterns encoded in protein 3D structures. Here, we present G-SPRI, a multilevel framework built on a novel alpha-shape protein graph that accurately captures residue connectivity from atomic-resolution geometry and enables precise message passing around mutation sites. Using this graph representation, G-SPRI integrates wild-type structural properties and mutation-specific perturbation signals derived from the Protein Data Bank (PDB) universe to support graph-based learning for distinguishing pathogenic from benign missense variants. G-SPRI performs strongly across multiple key tasks. On the binary prediction benchmark, G-SPRI delivers improved pathogenicity prediction for individual mutations. By integrating mutation recurrence across the pan-cancer cohort, G-SPRI recovers more known cancer driver genes than state-of-the-art methods from more than 2.3 million mutations. Furthermore, by jointly quantifying site-specific pathogenicity and co-clustering influence within higher-order structural organization units, G-SPRI provides comprehensive evidence for pinpointing likely driver mutations and structurally susceptible regions within disease genes.

## INTRODUCTION

Missense mutations are among the most common mutation types in the coding regions of the human genome^1–3^. While many missense mutations alter protein functions and contribute to disease progression^4–6^, a large fraction are presumed to be functionally neutral^7^. Identifying disease-causing missense mutations from this background is therefore a critical task that can guide downstream analyses, including the identification of genotypes underlying Mendelian disorders and the development of mutation-specific targeted therapies for complex human diseases such as cancer^8–12^.

Previous studies have primarily relied on expert-curated features or machine-generated embeddings derived from sequence information (e.g., evolutionary profiles, protein language models, co-evolution patterns, meta-analysis, and population genetics)^13–19^. However, these approaches often overlook the rich biophysical features encoded in protein three-dimensional (3D) structures and provide limited interpretability regarding the underlying biological mechanisms. The rapid growth of experimentally determined structures in the Protein Data Bank (PDB)^20^, together with increasingly accurate computationally predicted structures^21,22^, now enables broader integration of structural information into mutation-effect prediction. Several structure-centric methods, such as DAMpred, Rhapsody, and SPRI^23–27^, have been developed to characterize biophysical and dynamic properties using atomic-resolution information. These methods not only deliver comparable or superior accuracy but also enhance mechanistic understanding of mutations while potentially reducing computational cost.

However, existing methods face two key limitations. First, protein function often depends on the collective contribution of multiple residues within a contiguous spatial domain or structural scaffold, such as the catalytic domain of an enzyme^28,29^. Characterizing the influence of the neighboring region surrounding a mutation site, therefore, complements the intrinsic properties of the mutation itself. Although some methods use coarse-grained connectivity (e.g., centroid graphs) or implicit proximity information (e.g., co-evolution, structural context, or LLM attention over linear context) to estimate such neighboring influence^13,14,18,23,30^, a precise framework for identifying spatially relevant residues within complex structural patterns and quantitatively assessing the impact of the local environment remains challenging.

The second limitation of existing methods is their limited effectiveness when analyzing real-world mutation profiles using only predicted computational pathogenicity scores for individual mutations. For example, ∼55% of somatic mutations in the publicly released TCGA dataset are annotated as likely damaging by PolyPhen-2^15^. Such results suggest a high false-positive rate and make it difficult to prioritize high-confidence cancer driver mutations. We attribute this limitation primarily to two factors. First, most existing methods do not comprehensively integrate site-specific pathogenicity at individual mutation sites, co-clustering influence in local structural regions, and cohort-level recurrence across patients. As a result, current methods suffer from both design and performance limitations, leading to high false-positive rates and reduced accuracy in pathogenic-variant prioritization. Second, there is limited validation of the large-scale mutation landscapes in real-world mutation profiles. In addition, data leakage during model training can arise when mutations from the same protein appear in both training and test sets, inadvertently leaking gene-level information^13,27^.

To overcome these limitations, we developed G-SPRI, a comprehensive framework for analyzing missense mutations across the human genome. G-SPRI constructs an alpha-shape protein graph that enables precise layered *k*-hop message passing based on accurate residue connectivity^31–34^. It computes wild-type structural context and mutation-induced perturbations estimated from the PDB universe as node attributes and leverages a graph attention network to predict the functional impact of individual mutations^24,34,35^. G-SPRI further enables whole-genome gene prioritization by integrating functional impact and mutation recurrence within a disease cohort. In addition, G-SPRI quantifies co-clustering influence and offers a unique capability to pinpoint cancer driver mutations and structurally susceptible regions. Together, these features provide several notable advantages. First, G-SPRI outperforms several state-of-the-art methods in the evaluated PDB-mappable benchmark setting, including AlphaMissense, EVE, gMVP, and PolyPhen-2^13,15,18,30^, on a synthetic dataset integrating PDB-mappable ClinVar variants and putative benign variants meticulously selected from real-world mutation profiles^12^. Second, in the full-coverage experimental PDB setting, it identifies more known cancer driver genes than Dig and MutPanning^36,37^ among the top-ranked whole-genome candidates. Third, it provides fine-grained predictions for disease-relevant mutation sites and susceptible structural domains, complementing gene-level prioritization. Finally, its structure-derived features capture essential biological mechanisms of mutations and their neighboring effects, allowing G-SPRI to operate effectively in a sequence-free setting (without MSA or LLM embeddings) and potentially enabling broader coverage of proteins for which MSA construction is challenging, such as disordered regions and orphan genes.

## RESULTS

### Alpha-shape protein graph construction and graph-based learning for mutation-effect prediction

As shown in **Figure 1A**, the framework begins by representing a protein structure as an alpha-shape protein graph, which provides a geometrically grounded description of residue connectivity^24,31–33,38–40^. Unlike previous graph formulations, such as *k*-nearest-neighbor (*K*NN) centroid graphs, voxel graphs, or co-evolution graphs^13,30,41,42^, our approach is designed to reduce spurious residue contacts and preserve layered spatial relationships among neighboring residues. Starting from the 3D atomic coordinates of a protein, we compute an atomic-resolution weighted Voronoi diagram using atom-specific van der Waals (VDW) radii, and then trim the resulting structure with an alpha value of 1.4 Å, corresponding to the approximate radius of a water molecule. This procedure defines atomic connectivity based on physically meaningful geometric proximity. The resulting atom-level graph is then mapped to the residue level, where nodes represent residues and edges represent residue pairs connected through their constituent atoms, yielding a residue-connectivity graph and its corresponding contact-map adjacency representation.

**Figure 1.**
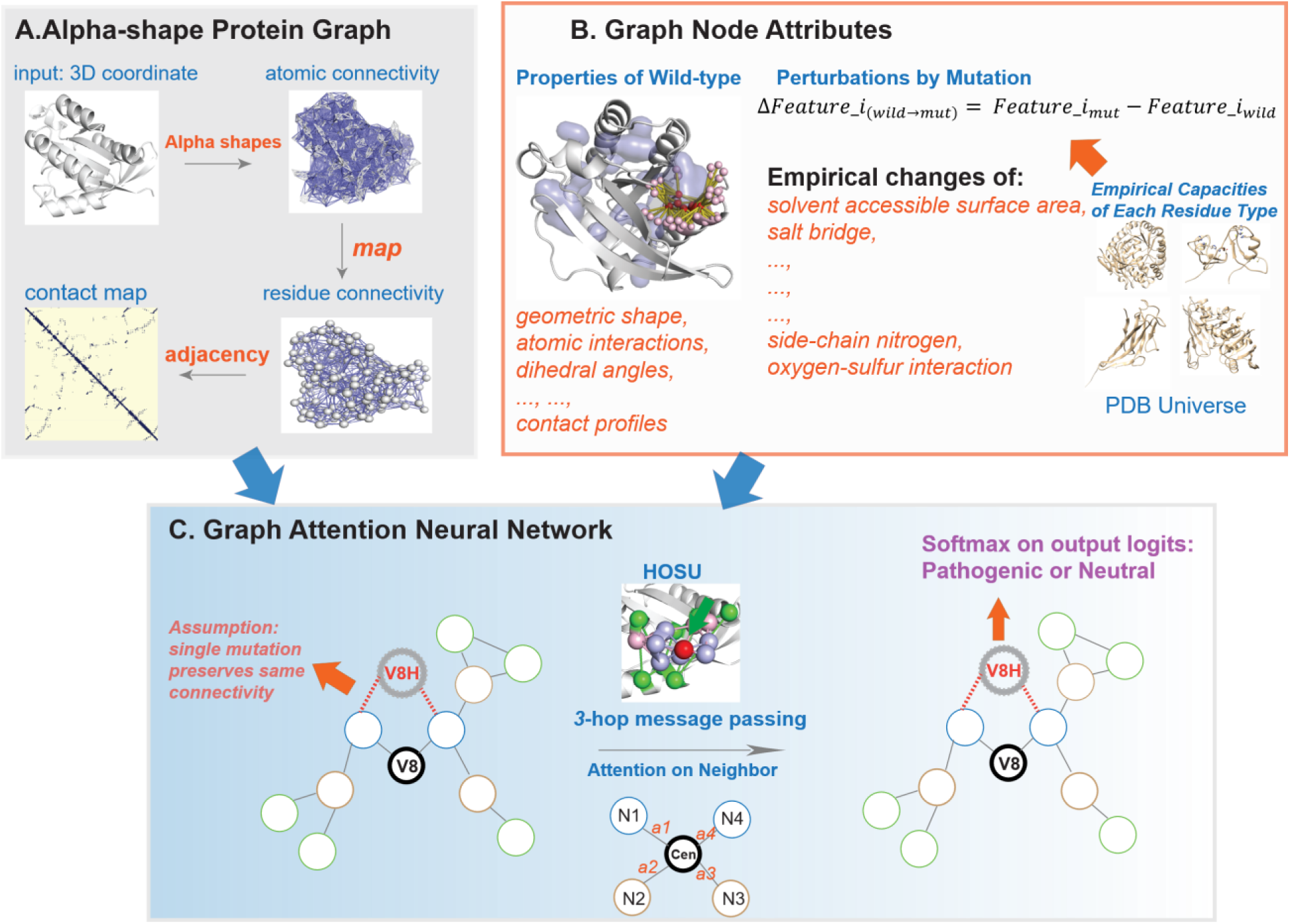
Architecture of G-SPRI for predicting the pathogenicity of individual missense mutations. **A**. G-SPRI constructs an alpha-shape protein graph. **B**. G-SPRI computes structural propensities of the wild-type residue in the reference conformation, and estimates the perturbation caused by the mutant residue type. **C**. G-SPRI assumes that a single missense mutation preserves the same graph connectivity. A graph attention network using 3-hop message passing is employed to distinguish pathogenic from benign variants.

Building on this graph representation, the second component of the framework defines graph node attributes that capture both the baseline structural context of the wild-type conformation and the perturbations introduced by the mutation (**Figure 1B**). For each residue node, wild-type features encode local structural properties and physicochemical constraints^24^, including geometric shape, atomic interactions, dihedral angles, salt bridges, and contact profiles. To quantify mutational changes, we further calculate feature perturbations as differences between mutant and wild-type states, thereby describing empirical changes induced by amino acid substitution. These perturbation features include changes in solvent-accessible surface area (SASA), salt bridges, side-chain nitrogen contacts, oxygen-sulfur interactions, and related structural descriptors^43–45^. In parallel, empirical capacity statistics for each residue type are derived from the PDB universe to provide a broader structural prior that contextualizes residue-specific behavior.

These graph connections and node attributes are then integrated into a graph attention neural network for pathogenicity prediction (**Figure 1C**)^34,35^. Under the assumption that a single amino acid substitution does not substantially alter the underlying residue connectivity, the mutant residue is introduced onto the same alpha-shape graph while its node features are updated to reflect the corresponding mutational perturbation. The network performs *k*-hop message passing, illustrated here as 3-hop propagation, to aggregate information from neighboring residues by exploiting the layered organization captured in higher-order structural organization units (HOSUs)^24^. Attention mechanisms further allow the model to weight neighboring residues according to their implicit structural relevance, thereby improving quantification of neighboring effects. Finally, the learned graph representation is passed to a softmax classifier that outputs the probability that a variant is pathogenic or benign.

Overall, this framework establishes a unified structure-centric learning pipeline in which the alpha-shape protein graph serves as the foundation for constructing biologically and geometrically meaningful residue connectivity, extracting mutation-aware structural features, and supporting precise graph-based inference. By more faithfully encoding spatial organization and layered residue proximity, this framework enables more accurate message passing and provides the basis for downstream analyses throughout the study, including contact-map generation, feature computation, and higher-order structural organization unit characterization.

### Development of G-SPRI for pathogenicity prediction of missense mutations

Building on these algorithmic innovations (**Figure 1**), we developed G-SPRI as a multilevel framework for systematic analysis of missense mutations in the human genome. The framework consists of three major components. At the variant level, G-SPRI predicts the functional impact of individual missense mutations using a structure-centric graph-learning strategy. At the gene level, it combines predicted pathogenicity scores with mutation recurrence in disease cohorts to compute gene-level risk scores and prioritize candidate genes across the genome. At the structural-region level, it quantifies co-clustering influence together with site-specific pathogenicity to separate potential driver mutations from passenger mutations within the same gene and to reveal regions of elevated structural susceptibility.

To support variant-level prediction, G-SPRI encodes each protein as an alpha-shape protein graph whose node attributes integrate both wild-type structural information and mutation-specific perturbation signals. One feature group describes the structure-derived properties of the wild-type residue reflecting biological constraints in its reference-state conformation, largely following the SPRI design^24^. The second feature group captures the expected alterations caused by amino acid substitution, estimated from structure-derived statistics for amino acid types collected from a non-redundant PDB universe^20^. In this way, G-SPRI combines reference-conformation context with mutation-specific change profiles within a unified graph representation.

The resulting features are vectorized and concatenated as node attributes to form the complete graph:

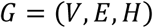

where *V* represents nodes, *E* represents edges, and *H* represents node attributes. Based on this graph formulation, G-SPRI uses a graph attention network to learn patterns that distinguish pathogenic from benign mutations. To minimize information leakage and ensure rigorous performance evaluation, training, validation, and test sets are separated using a leave-graph-out strategy based on protein-level stratification.

### Benchmarking binary classification of individual mutations

To assess the predictive performance of G-SPRI, we compared it with state-of-the-art approaches on a synthetic test dataset integrating ClinVar-PDB and putative benign variants meticulously selected from real-world mutation profiles^12^. The benchmark included AlphaMissense^13^, a structure-informed protein language model; EVE^18^, an evolutionary generative model; gMVP^30^, a co-evolution graph neural network; and PolyPhen-2^15^, a commonly used alignment-based predictor. Model performance was evaluated comprehensively using the area under the precision–recall curve (AUPRC) and the area under the receiver operating characteristic curve (AUROC), as summarized in **Figure 2**.

**Figure 2.**
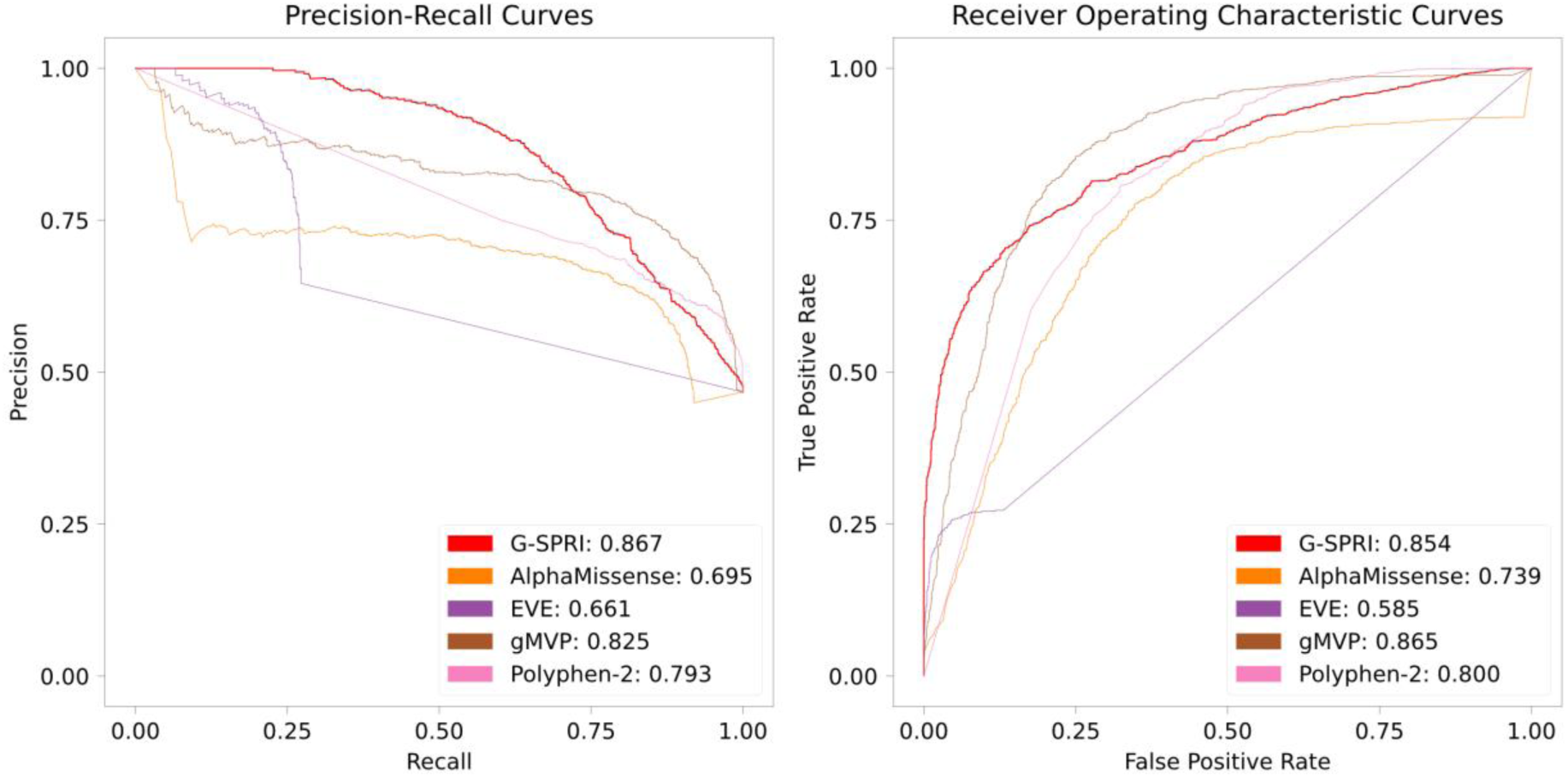
Precision-recall curves (PRC) and receiver operating characteristic curves (ROC) for one fold of the test dataset synthesized from ClinVar and putative benign variants. Note that the values are based on one fold of the test dataset. We also report average values of five test folds in the main text.

These results demonstrate that G-SPRI achieved strong overall performance for both the AUPRC and AUROC. Across five test folds, G-SPRI achieves the highest mean AUPRC of 0.827, exceeding AlphaMissense (0.713), EVE (0.756), gMVP (0.822), and PolyPhen-2 (0.807). A similar trend is observed in the receiver operating characteristic curve, where G-SPRI exhibits the second highest mean AUROC of 0.804, superior to AlphaMissense (0.755), EVE (0.646), and PolyPhen-2 (0.797), while gMVP achieves the highest AUROC of 0.842. We also reported the values based on one fold of test dataset in **Figure 2**. We emphasize that benchmark performance alone is insufficient to establish practical utility, because the TCGA pan-cancer mutation profiles contain more than 200 times as many missense mutations as these curated benchmark datasets.

### G-SPRI enables whole-genome gene prioritization in the TCGA pan-cancer cohort

Building on the accurate prediction of pathogenicity for individual missense mutations, we next extended G-SPRI to analyze mutation profiles at the disease-cohort level. As illustrated in **Figure 3A, B**, G-SPRI first estimates the pathogenic likelihood of each mutation and then integrates these site-specific predictions with mutation recurrence across patients within a cohort. This combination converts variant-level pathogenicity signals into quantitative gene-level risk scores, providing a principled basis for whole-genome gene ranking in a large cancer dataset.

**Figure 3.**
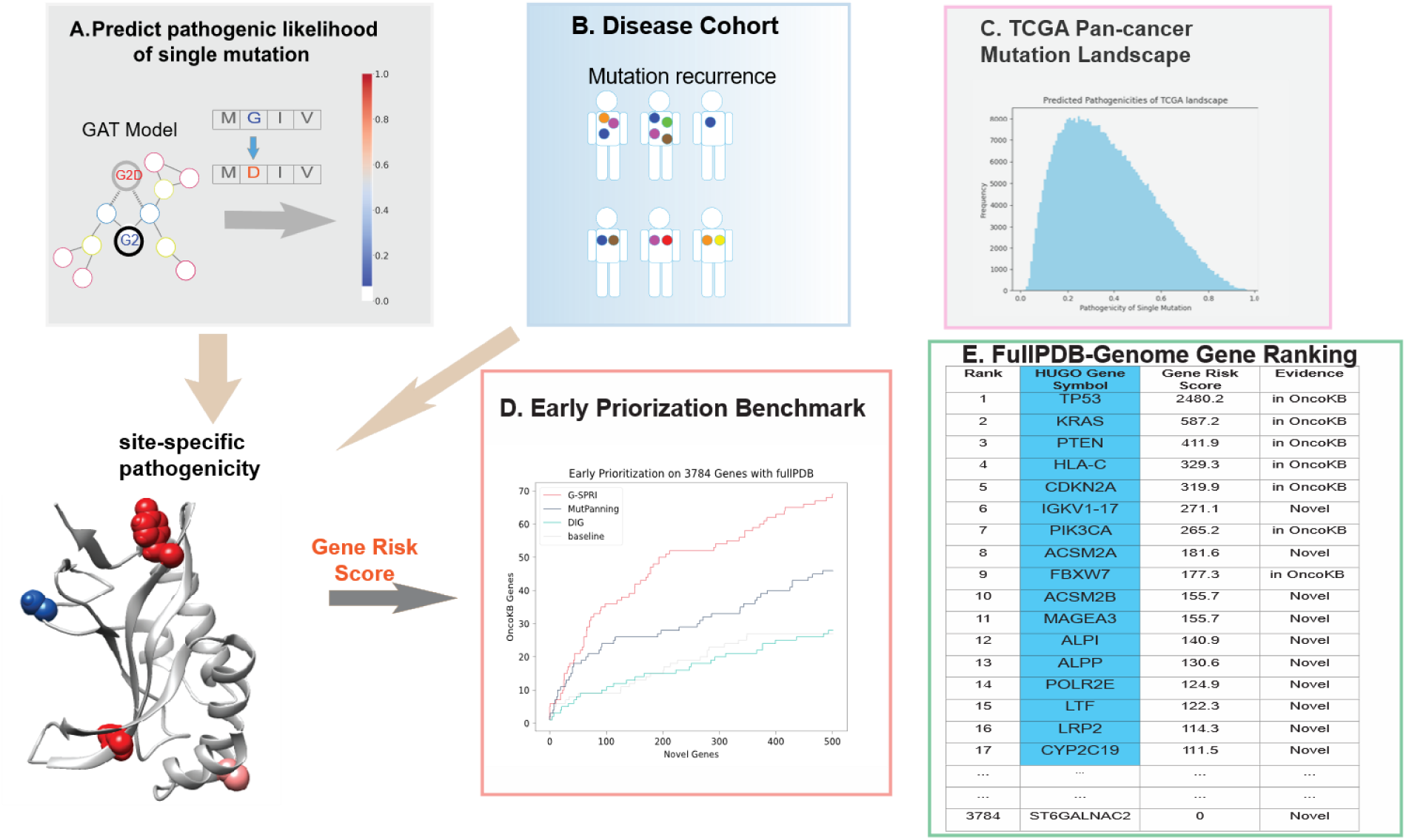
G-SPRI integrates pathogenicity prediction of individual mutations and recurrence within the disease cohort to compute gene risk scores, and systematically ranks disease-relevant genes across the whole human genome. **A**. G-SPRI computes pathogenic likelihood of all observed missense mutations in the disease cohort. **B**. Mutation recurrences are extracted from patients’ mutation profiles. **C.** G-SPRI reveals a plausible predicted mutation landscape in which the majority of mutations have minor or moderate impacts. **D**. G-SPRI exhibits improved performance in identifying known cancer driver genes using an early prioritization benchmark. **E**. G-SPRI provides gene rankings at the-genome scale, we illustrate ranking at fullPDB-genome scale.

This cohort-level framework was evaluated in the TCGA pan-cancer dataset to determine whether G-SPRI can recover known cancer driver genes from real-world mutation landscapes^10^. Following a strategy similar to MutPanning^37^, we analyzed more than 2.3 million missense mutations classified as somatic or private germline variants. The predicted pathogenicity distribution across the TCGA pan-cancer landscape is shown in **Figure 3C** and displays a plausible global pattern, with 46.1% of mutations falling in the confidently benign probability range (p < 0.3) and 78.1% of mutations falling into in benign or likely benign ranges (p < 0.5). These model-based categories should be interpreted as prioritization labels rather than definitive functional annotations, but the distribution is consistent with the expectation that most observed mutations in cancer genomes are passengers rather than functional drivers.

To assess gene-ranking performance, we next applied two complementary evaluation strategies. The first was an early-prioritization analysis, which measures the enrichment of known cancer driver genes among the top-ranked candidates^36^. The second was a biological interpretation of novel genes prioritized by G-SPRI, focusing on whether these candidates have independent functional or disease-related supporting evidence. Together, these analyses evaluate not only ranking performance but also the biological relevance of newly predicted genes.

This benchmark revealed the clearest advantage of G-SPRI in the subset of genes with full-coverage experimental PDB structures, where structural information is most reliable. Among 3,784 such genes, 208 are annotated as cancer driver genes in OncoKB with support from at least two sources^10^. In this high-confidence setting, G-SPRI identified 36, 50, and 69 known cancer genes when the top-ranked background gene set reached 100, 200, and 500 genes, respectively, whereas MutPanning identified 24, 28, and 46 genes at the same thresholds^37^, and Dig showed performance similar to the recurrence rate-only baseline^36^. As further illustrated by the early-prioritization curves in **Figure 3D**, G-SPRI consistently outperformed MutPanning, Dig, and the recurrence-only baseline, indicating that integration of site-specific pathogenicity with recurrence provides a substantial advantage in recovering established cancer drivers from a noisy mutational background. Because this subset has the highest-quality structural coverage, it also provides the fairest setting for direct comparison.

This performance advantage became less pronounced when the analysis was expanded beyond full-coverage PDB structures. For genes with acceptable-quality PDB structures, including both full-coverage and partial-coverage cases, G-SPRI performed on par with MutPanning, and recovered more known genes than Dig and the baseline model. The early prioritization performance is illustrated in the Supplemental Information. We also evaluated genes represented by AlphaFold2-predicted structures in the Supplementary Information, but these results should be interpreted cautiously because structural incompleteness or modeling inaccuracies may introduce additional uncertainty into pathogenicity prediction and downstream gene ranking; similar inconsistencies in protein-property prediction have also been reported^46^.

These benchmarking results are further supported by the ful<u>l</u> PDB-mappable genome ranking output generated by G-SPRI. As shown in **Figure 3E**, top-ranked genes include well-established cancer drivers such as TP53, KRAS, PTEN, CDKN2A, and PIK3CA, as well as additional genes with literature support for cancer relevance including FBXW7, MAGEA3, LTF, ALPP, ACSM2B, ALPI, LRP2, CYP2C19, and POLR2E. For example, FBXW7 encodes the substrate-recognition component of the SCF ubiquitin ligase and functions as a tumor suppressor based on experimental evidence that cancer-associated hCDC4/FBXW7 mutation disrupted cell-cycle regulation of cyclin E, while reintroduction of hCDC4 restored periodic cyclin E expression in a breast carcinoma-derived cell line, supporting FBXW7 loss as a mechanism of oncogenic cell-cycle deregulation^47^. MAGEA3, a cancer-testis antigen, has also been functionally implicated in tumor progression: in cervical cancer models, MAGE-A3 knockdown suppressed proliferation, migration, invasion, EMT/Wnt signaling and tumor growth, whereas MAGE-A3 overexpression produced the opposite phenotype^48^. LTF encodes lactotransferrin/lactoferrin and has tumor-suppressive activity in nasopharyngeal carcinoma, where LTF represses AKT signaling by suppressing PDK1 and interfering with the K18–14-3-3 complex, thereby inhibiting NPC tumorigenesis^49^. ALPP, encoding placental alkaline phosphatase, is an actionable cancer-associated surface antigen supported by experimental evidence: EGFR inhibition in lung adenocarcinoma induced ALPP through FoxO3a activation, and combined EGFR TKI plus ALPP–MMAF antibody-drug conjugate enhanced tumor killing in osimertinib-sensitive and –resistant lung cancer models^50^. ACSM2B has been reported as a lipid-metabolism enzyme and potential negative regulator of liver cancer progression^51^. ALPI is a differentiation-associated colorectal cancer marker, as intestinal alkaline phosphatase was induced by HDAC inhibitors in colon cancer cells through a KLF5-dependent mechanism^52^. CYP2C19 downregulation was associated with aggressive hepatocellular carcinoma and independently predicted poorer recurrence-free survival, supporting its use as a cancer-associated prognostic gene rather than a mechanistic driver^53^. LRP2 was linked to cancer dedifferentiation through epigenetic silencing across multiple solid tumor types and low expression was associated with poorer outcome^54^. POLR2E has cancer-susceptibility evidence through the rs3787016 polymorphism, which was associated with overall cancer and prostate cancer risk in a case-control/meta-analysis^55^. Together, these findings suggest that G-SPRI is capable of recovering canonical driver genes and prioritizing previously underappreciated genes that may warrant further biological investigation. Collectively, the biological plausibility and supporting evidence for these candidates further support the utility of G-SPRI as a hypothesis-generating tool for genome-wide cancer gene discovery.

### G-SPRI pinpoints driver mutations and susceptible regions through co-clustering influence

Building on gene-level prioritization, G-SPRI further refines mutation interpretation by integrating site-specific pathogenicity with co-clustering influence at the structural-region level. This combined framework is designed to distinguish likely driver mutations from passenger mutations occurring within the same gene while also highlighting HOSUs that are especially susceptible to disease-associated variation and may indicate functional domains. By combining the intrinsic pathogenic potential of a mutation site with the pathogenic context of nearby clustered mutations, G-SPRI moves beyond gene-level signals to achieve more precise, site-resolved interpretation.

Figure 4 uses PTEN as a representative example to illustrate how this integrated scoring strategy prioritizes mutation sites. As shown in the upper panel, G-SPRI ranks PTEN mutation sites by combining site-specific pathogenicity and co-clustering influence into a unified score. Among 135 residue sites that harbor mutations in pan-cancer cohorts, G-SPRI identifies R130 as the <u>most</u> top-ranked site, reflecting exceptionally strong site-specific pathogenicity together with an additional contribution from neighboring co-clustering influence. Other highly ranked sites, including D92 and R173, also show substantial contributions from both components, supporting their prioritization as functionally important mutation hotspots. In contrast, low-ranked sites such as D312 exhibit minimal site-specific pathogenicity and little co-clustering support, consistent with a limited contribution to disease relevance. Beyond mutation sites with clear pathogenicity patterns, it is challenging to distinguish between G36 and R14, which exhibit similar site-specific pathogenicity. After co-clustering influence is quantified, G36 shows stronger co-clustering influence than R14 and receives a higher unified-score ranking (13 vs 40).

**Figure 4.**
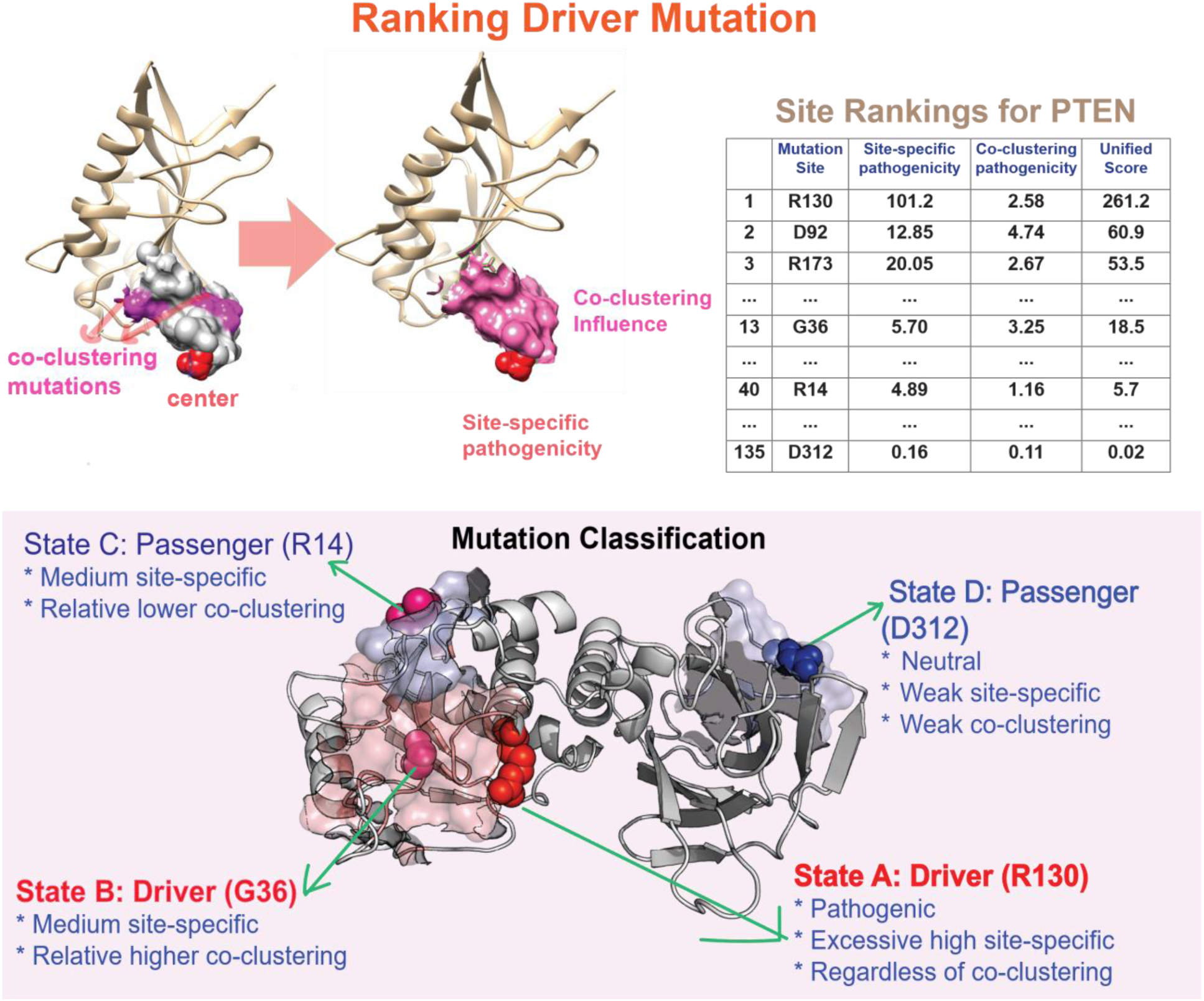
G-SPRI quantifies co-clustering influence, together with site-specific pathogenicity of the center residue, which provides a unique capability to reveal granulated details, including distinguishing driver mutations from passenger mutations occurring in the same gene. PTEN is used here as a representative example.

This integrated ranking naturally leads to a classification framework that separates mutations into distinct functional states. As illustrated in the lower panel of Figure 4, State A represents driver mutations with pathogenic effects driven primarily by exceptionally high site-specific pathogenicity, leading to a high unified score regardless of co-clustering strength. State B also represents driver mutations, but in this case, the pathogenic interpretation arises from the combination of moderate site-specific pathogenicity and strong co-clustering influence. By contrast, State C includes pathogenic but likely passenger mutations that show only moderate site-specific pathogenicity and weak co-clustering, whereas State D corresponds to neutral passenger mutations lacking both pathogenicity and co-clustering evidence. It is difficult for site-specific evidence alone to distinguish mutation sites assigned to States B or C, whereas G-SPRI provides a comprehensive structural framework for distinguishing mutations that are individually dominant from those whose importance depends on their pathogenic neighborhood.

Taken together, Figure 4 shows that G-SPRI can extend gene-level prioritization to structural interpretation of mutation sites and regions. By jointly modeling site-specific pathogenicity and co-clustering influence, the framework not only helps distinguish candidate driver mutations from likely passenger mutations within the same gene, but also identifies structurally vulnerable regions that may concentrate disease-associated effects. This capacity is particularly important for mutational hotspot genes such as PTEN, where multiple mutations occur across the structure but differ substantially in their functional and clinical significance.

## Discussion

In this study, we developed G-SPRI as a multilevel framework for systematic analysis of missense mutations, spanning variant-level pathogenicity prediction, cohort-level gene prioritization, and site-resolved identification of driver mutations and susceptible structural regions. G-SPRI offers several advantages for predicting disease-relevant mutations. First, it formulates an alpha-shape protein graph that precisely captures residue connections in proteins, thereby improving feature aggregation and layered *k*-hop message passing in graph neural network models; Second, it computes structural, biophysical, and geometric features that reflect biological functions encoded in protein structures, improve performance on the binary prediction task, enhance interpretations of the graph model, and potentially increase prediction coverage for mutations in disordered regions or orphan genes because it does not require evolutionary signals from MSA; Third, by integrating predicted pathogenicity for individual mutations with mutation recurrence within disease cohorts, G-SPRI exhibits improved early prioritization performance in identifying cancer driver genes in pan-cancer data, with supporting evidence for top-ranked novel candidate genes; Finally, G-SPRI can distinguishes driver mutations from passenger mutations within the same gene and identifies high-susceptibility structural regions by quantifying co-clustering influence with site-specific pathogenicity.

Despite these strengths, several limitations should be considered. First, the performance of G-SPRI depends on the availability and quality of protein structures, and the clearest advantage in gene prioritization was observed for genes with full-coverage experimental PDB structures. Although acceptable-quality partial structures and predicted models can expand coverage, inaccuracies or incompleteness in these representations may reduce robustness in downstream predictions. We expect this limitation to be alleviated by the growing availability of PDB structures and by continued improvement in structure-prediction methods. Second, the current framework assumes that a single amino acid substitution does not substantially alter the underlying residue connectivity, which is a practical approximation for graph construction but may be less accurate for strongly destabilizing mutations or conformationally disruptive variants. Third, the G-SPRI framework does not consider the impact of loss of heterozygosity (LOH) when calculating gene-level risk scores. Fourth, the perturbation features inferred from the PDB universe provide an efficient approximation of mutation-induced effects, but they do not fully replace direct structural remodeling or explicit energetic simulation of each mutant protein. Fifth, the current evaluation focuses primarily on missense mutations, and the generalizability of the framework to other variant classes, such as indels, remains to be established.

Additional limitations arise at the cohort-analysis level. The gene-prioritization framework integrates pathogenicity with recurrence, but cancer driver discovery is also influenced by tumor type, mutational impact on specific function mechanisms, selective pressures, tumor-type-specific background mutation processes, and noncoding, copy-number, or LOH events that are not explicitly modeled here. Similarly, the co-clustering framework provides a useful structural interpretation of localized mutational burden, but the biological significance of newly prioritized hotspots and HOSUs will require further experimental and clinical validation. Future work could therefore extend G-SPRI by incorporating improved structure sources, mutant-specific structural refinement, cancer-type-specific modeling, calibration analysis, and multimodal genomic evidence. Such developments would further strengthen the ability of structure-based learning frameworks to support mechanistic interpretation and translational discovery in human disease. Such developments would further strengthen the ability of structure-<u>centric</u> learning frameworks to support mechanistic interpretation and translational discovery in human disease.

## Acknowledgments

The resources from the Quantitative Biomedical Research Center (QBRC) and BioHPC at UT Southwestern Medical Center are gratefully acknowledged.

## Funding

This work was supported by the following funding: the Rally Foundation, Children’s Cancer Fund (Dallas), the HHOW Award, the Sam Day Foundation Award, the Cancer Prevention and Research Institute of Texas (RP180319, RP200103, RP220032, RP170152 and RP180805), and the National Institutes of Health funds (R01DK127037, R01CA263079, R21CA259771, R21CA273282, P30CA142543, UM1HG011996, R01NS142141, R01CA284591, R01DK130961, and R01HL144969) (to L.X.).

## Contributions

BW and LX conceived and designed the study. BW, AF, and BY developed the graph neural network and performed the benchmarking. LY and ZL collected the TCGA raw data. BW and LY annotated and curated TCGA mutation profiles. LX acquired the funding. BW and LX wrote the manuscript. All co-authors have revised and approved the final manuscript.

## Corresponding authors

Correspondence to Lin Xu.

## Ethics declarations

The authors declare that they have no competing interests.

## Methods

### Alpha-shape Graph Computation

In a typical protein graph (contact map), residues are represented as nodes, with edges denoting residue-residue connections. Existing methods for generating protein graphs can generally be classified into three categories: centroid graphs, co-evolution graphs, and voxel graphs^13,14,18,23,30^. However, the simplicity of these approaches may introduce inaccurate connections, leading to suboptimal feature aggregation during message passing in a graph neural network. A more detailed discussion of the limitations of these graph representations is provided in SI Section 1.1.

To overcome the limitations of existing protein-graph representations, we introduce a novel alpha-shape protein graph in this study. Alpha-shape computation for proteins is a well-established approach for analyzing 3D structures. It enables characterization of geometric features (pockets, surface, and buried regions), calculation of solvent-accessible surface area (SASA), identification of protein-protein interaction (PPI) regions, and extraction of atomic interactions and residue-level contact profiles with layered relationships^31–33,38–40,44,45^. Following standard practices in protein-graph construction, our approach derives residue-level connectivity from a given high-quality 3D structure. Specifically, we computed alpha shapes for each protein structure at an alpha value of 1.4 Å, corresponding to the radius of a water molecule. This process corresponds to weighted Delaunay triangulation (the mathematical dual of the Voronoi diagram), incorporating the van der Waals (VDW) radii of each atom type and trimming simplices based on the selected alpha value. The resulting alpha shape provides accurate atomic connectivity, which we subsequently map onto a residue-level graph to define edges and nodes. The alpha-shape protein graph offers two key advantages over the commonly used centroid graph: (i) improved accuracy in identifying neighboring residues at short distances and (ii) the ability to precisely capture layered structural information at moderate distances regardless of local packing density, thereby enabling more orderly feature aggregation through *k*-hop message passing in graph neural networks.

We used the protein Rab-33B (PDB: 6ZAY, chain: A) as a case study to demonstrate the advantages of our approach. To identify residues connected to the center residue Gly147 within a short distance, the alpha-shape graph successfully captured neighboring residues Leu145 and Leu174 that were overlooked by the centroid graph. Conversely, the alpha-shape graph excluded Val113, Cys150, and Ala163, which were incorrectly inferred as neighbors by the centroid graph (see SI Section 1.2). This case highlights how the alpha-shape graph improves accuracy by eliminating spurious connections, thereby reducing noise and preventing erroneous message passing in graph neural network models. In addition, when identifying neighboring residues at moderate distances, the alpha-shape graph revealed layered relationships that the *k*-NN centroid graph failed to capture (see SI Section 1.3). This layered organization further enables *k*-hop message passing with improved precision in feature aggregation.

The alpha-shape protein graph was computed using the in-house C source code. The alpha-shape graph captures atomic interactions for both intra-residue (within the same residue) and inter-residue (between different residues). The inter-residue atomic interactions were used to construct the residue-level protein graph (contact map). For each graph, the connectivity information is stored as an undirected (bidirectional) adjacency list and adjacency matrix.

### Node Attributes Computation

G-SPRI derives features from two sources: the structural properties of the wild-type site in the reference conformation and the perturbation of these structure-derived features across residue types. We applied SeqMapPDB to identify genes in ClinVar with mappable PDB structures and largely followed SPRI to compute structural information for the wild-type residue. We then used MMseqs2 to generate a non-redundant PDB universe with sequence identity below 50% for sequence clustering. After quality control, 28,321 PDB chains were used to calculate structure-derived properties for each of the 20 amino acid types. To reflect the different capacities of amino acids, we selected the maximum, 99th percentile, and mean values of the features for each amino acid type and estimated empirical perturbation values associated with each mutant residue type. For occasional missing features, we applied *k*-NN imputation. We then performed min-max normalization to obtain normalized values.

### GNN Training Regime

We used PyTorch Geometric (PyG) and PyTorch to implement the graph attention network. We chose *k* = 3 convolution layers (3-hop), which capture information from nodes up to three layers away from the mutation site; the channel sizes for the 1-hop, 2-hop, and 3-hop layers were set to 128, 512, and 1, respectively. We used ReLU as the activation function, Adam as the optimizer, a fixed random seed, a learning rate of 0.0001, and weight decay of 0.0005.

In addition to the known benign and pathogenic mutations annotated by ClinVar, we also considered the wild-type residue as benign in the training dataset, with perturbation features assigned a value of 0 to maintain a consistent feature dimension. We employed binary cross-entropy (BCE) as the loss function. Early stopping was applied when there was no improvement in AUROC and AUPRC on the validation dataset, with a patience of 500 epochs. We selected the epoch with the highest combined AUROC and AUPRC on the validation set rather than the training set to reduce overfitting. We used grid search to select the optimal hyperparameters, including hidden channels, dropout rate, the positive-class weight in the loss function, learning rate, and weight decay. Detailed hyperparameter ranges are provided in the **Supplementary Information**.

We applied the softmax function to the output logits to obtain the probability of functional impact for each mutation, ranging from 0 to 1, recorded as *D*_*mut*_. A lower probability indicates a lower likelihood of a damaging effect, and vice versa. G-SPRI captures essential distinguishing features as the initial inputs and demonstrates strong computational efficiency and feasibility; a typical training run for a single fold took ∼28 minutes on a workstation equipped with a single Nvidia RTX-3090Ti GPU and an AMD Ryzen 7900X CPU.

### Benchmarking Dataset

To guide model learning and support a fair comparison, we integrated ClinVar and putative benign variants meticulously selected from real-world mutation profiles as the benchmark dataset. We selected genes with high-quality PDB structures and designated the resulting set as ClinVar-PDB. For the ClinVar-PDB dataset, we downloaded the VCF file from the ClinVar FTP site. Because ClinVar does not provide reference protein IDs, we used the Ensembl VEP online annotation tool (v114, GRCh38) to obtain NCBI RefSeq transcript and protein IDs. Reference sequences for each protein were downloaded from NCBI. Mutation-site information was cross-checked against the corresponding sequences, and a small number of proteins with mismatched mutation records were excluded. Because VEP may provide multiple transcripts or protein isoforms for a single gene, we selected a single protein isoform per gene to avoid duplication. We then used the SeqMapPDB pipeline to obtain PDB-mappable proteins. As a result, 1,844 proteins were fully covered, and 532 were partially covered. In total, we collected 5,432 benign and 14,865 pathogenic mutations mapped across 2,533 PDB chains (graphs). For putative benign variants, we focused on genes with available PDB structures, removed any OncoKB genes, and narrowed the selection to genes without pathogenic mutation records in ClinVar. We further selected genes without hotspot mutations (> 3 recurrences in TCGA pan-cancer), and with missense mutation rates lower than the median across the human genome. As a result, an additional 4,394 putative benign mutations from 201 genes were included in the benchmarking.

### ***k-***fold Leave-Graph-Out Cross Validation

Conventional methods typically use *k*-fold cross-validation to evaluate model performance. However, this approach may lead to data leakage, because mutations from the same gene or protein may appear in both training and test datasets and thus leak gene-level information into model development. To address this limitation, we employed a modified *k*-fold leave-graph-out cross-validation scheme, which uses mutually exclusive graphs (roughly representing proteins or genes) across the training, validation, and test datasets.

For the benchmark dataset, we set *k* to 5. Each fold includes 250 ClinVar-PDB graphs for validation, 250 ClinVar-PDB graphs and randomly splitting putative-benign protein graphs for testing, and the remaining ClinVar-PDB graphs for training. To obtain fair comparisons with other methods (AlphaMissense, EVE, gMVP, PolyPhen-2), we downloaded or obtained prediction results from their websites or databases. A pairwise alignment was employed to address potential sequence index shifts between different protein isoforms used.

### Gene Risk Score and Genome-wide Ranking

To analyze real-world mutation profiles for a given disease cohort, we further extended the use of predicted pathogenicity scores by integrating mutation recurrence across the disease cohort. For the cohort under investigation, we retrieved candidate missense mutations from individual patients. We then counted the recurrence of each mutation by iteratively searching all mutation profiles within the cohort. The graph model provided a damaging score, representing the likelihood of a pathogenic outcome for each mutation, on a probability scale from 0 to 1.

Since each wild-type residue can have 19 substitution patterns, we calculated the site-specific pathogenicity for a given mutation site as:

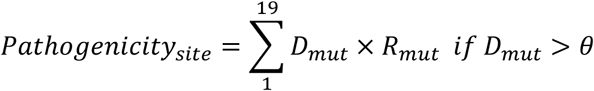

where the threshold for confidently classifying pathogenic impacts is adjustable for a specific disease type. In the TCGA pan-cancer cohort, it was set as 0.74.

We then summarized all the site-specific pathogenicity values across all sites within a protein to compute the gene-level scores, including the gene risk score and the normalized score.

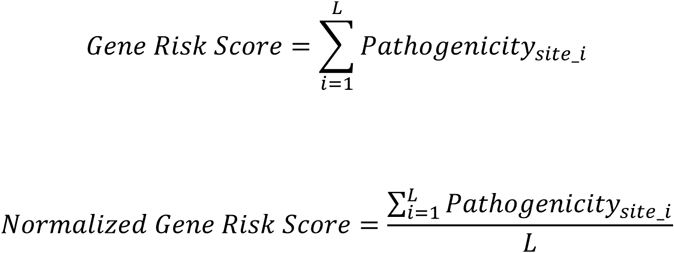

where *L* denotes the protein length.

In this way, we obtained gene-level risk scores in the human genome and gene rankings accordingly. Genes were ranked primarily by the gene risk score, while retained genes were also required to fall within the top 30% of normalized scores among all genes.

### TCGA Mutation Profiles

Following a strategy similar to MutPanning, we considered somatic variants as well as likely personalized germline variants. We retrieved GRCh37 DNA-variant records for somatic and personalized germline variants from the TCGA database and used a standalone Ensembl VEP tool (v115.1, GRCh37) to annotate them. Because genomic coordinate information may refer to multiple transcript or protein isoforms, we selected a single protein isoform per gene to avoid duplication; most of the selected isoforms were canonical. We then used gnomAD v4.1 to remove variants with a population minor allele frequency (MAF) greater than or equal to 0.001. gnomAD GRCh38 coordinates were converted to GRCh37 using liftOver. We identified ∼2.53 million unique missense mutations and counted their recurrence across the pan-cancer cohort. We also employed SeqMapPDB to map 3,784 genes to their full-coverage PDB structures, and map 1,498 genes to partial-coverage PDB structures. The remaining genes were included AlphaFold-2 v4 predicted structures. After additional quality control, including removal of cases with unaligned wild-type residues between VEP and the UniProt 2025-03 reference proteome and likely sequencing artifacts or annotated errors, pathogenicity was calculated for ∼2.3 million missense mutations.

For benchmarking with MutPanning and Dig, because they require synonymous mutations to model the background mutation rate, we input both missense and synonymous DNA variant information in their designated formats to obtain predictions.

### Co-clustering Mutation Influence

As a linear sequence folds into its 3D conformation, a spatial region often consists of residues with widely separated sequence positions, making mutation-enrichment patterns difficult to capture using linear sequence information alone. Existing methods for analyzing structural enrichment of mutations have several limitations, including (i) intuitive definitions of local spatial regions without consideration of structural arrangement (e.g., *k*-NN nearest-neighbor methods ignore local packing density) and (ii) enrichment metrics that frequently rely on mutation recurrence alone without additional quantification, such as functional impact. SPRI addressed these issues by integrating functional impact with mutation sites mapped to precisely defined spatial regions, demonstrating the ability to identify novel cancer driver mutations with low recurrence in pan-cancer cohorts. In this study, we further extended SPRI by developing a comprehensive metric of co-clustering influence.

G-SPRI has previously shown that integrating site-specific pathogenicity with mutation recurrence in a disease cohort effectively prioritizes cancer driver genes at the whole-genome scale. We further developed a co-clustering influence metric to quantify mutation enrichment in spatial regions, thereby pinpointing mutation sites and local structural regions with strong evidence of disease association. For a given mutation site (the center residue), co-clustering mutations are defined as neighboring sites that harbor mutations from other patients in the same disease cohort. Because protein function typically depends on residues in spatially proximal regions, it is reasonable to assume that these co-clustering mutations are more likely to perturb the same function and lead to similar phenotypic outcomes as the center mutation site. To overcome the limited availability of known functional domains, we constructed the HOSU for the center mutation site to represent a putative shared functional region and assumed that co-clustering mutations within the HOSU are more likely to affect the same biological function as the center site.

G-SPRI extends the SPRI framework for computing co-clustering mutation influence, reflecting the susceptibility of a local structural region. For each HOSU, we calculated site-specific pathogenicity scores for all neighboring mutation sites within the region and summarized their contributions. To account for varying local packing density, we further considered the total number of residues in the HOSU as a normalization factor and defined co-clustering mutation influence as follows:

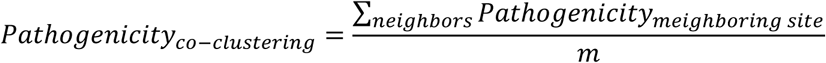

We then computed the combined evidence for each mutation site, defined as

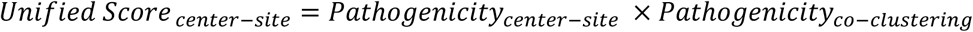

We then ranked mutation sites within the same gene according to this combined evidence *Unified Score*. Top-ranked mutation sites (top 20% used in this study) were classified as driver mutations and may show either (A) strong site-specific pathogenicity together with moderate or strong co-clustering influence or (B) moderate site-specific pathogenicity together with relatively strong co-clustering influence. Lower-ranked mutation sites may show either (C) moderate site-specific pathogenicity but weak co-clustering influence or (D) low site-specific pathogenicity regardless of co-clustering influence.

## Data availability

All data used in this study are publicly available and have been previously published; full details and citations are provided within the article.

## Code availability

https://github.com/Lin-Xu-lab/G-SPRI

## Notes

https://github.com/Lin-Xu-lab/G-SPRI

### Competing Interest Statement

The authors have declared no competing interest.

