## Supplementary material for "G-SPRI: A Structure-Centric Graph Model for Comprehensive Prediction of Cancer Driver Events from Missense Mutations": SI

### Supplemental Information

#### 1. Graph Representation of Proteins

##### 1.1 Brief Description and Limitations of Existing Protein Graph Representation Methods

Previous methods for generating protein graphs can be classified into three categories: centroid graphs, co-evolution graphs, and voxel graphs. The centroid graph considers only the centroid (typically the alpha- or beta-carbon atom) of each residue and uses a Euclidean distance threshold or a *k*-NN clustering to define connected residues (edges). However, it ignores complex spatial patterns, including local packing density, folding arrangements, and geometric orientations formed by other backbone and side-chain atoms. As a result, it may introduce erroneous connections, leading to imprecise feature aggregation during message passing in graph neural networks. In addition, centroid graphs do not clearly encode relative distances (i.e., layer/hop relationships) between residue pairs. Despite these limitations, they remain the most widely used graph representation for defining residue-level protein connections [1, 2, 3, 4].

Co-evolution graphs are pseudo-structural graphs that assume residues with co-evolutionary relationships also share spatial proximity and functional relevance [5]. However, co-evolution-derived connections may be unrealistic because they are inferred from statistical analyses of MSAs and additional assumptions (e.g., predefined lengths of flanking regions in protein contexts) [5, 6, 7]. In addition, this representation does not directly capture structural properties.

The voxel graph uses volumetric pixels to partition the 3D space of protein structures and can provide both atomic-resolution and residue-level information [8]. However, because atom types have different radii, fixed-size cubic voxels cannot fully represent spherical atoms.

##### 1.2 Comparison of the Alpha-shape Protein Graph and the Centroid Graph at Short Range

To demonstrate the advantages of the alpha-shape protein graph over the centroid graph, we used the Ras-related protein Rab-33B (PDB: 6ZAY, chain A; RefSeq: NP_112586.1) as a case study.

First, we examined the accuracy of connectivity at short range. We selected 147GLY as the center residue. In the centroid graph, a neighboring residue is considered connected to 147GLY if the distance between their centroids (beta carbons) is no greater than 7 Å. In the alpha-shape protein graph, residues are defined as connected to 147GLY if they share atomic interactions (first layer, i.e., 1-hop) with 147GLY. Detailed illustrations are provided in **Supplementary Figures 1-4**. These results show that the alpha-shape graph is more accurate than the centroid graph for identifying connected residues. The centroid graph incorrectly assigns edges between the center residue and 113VAL, 150CYS, and 163ALA, even though none of these residues share atomic interactions with 147GLY. In addition, the alpha-shape graph correctly captures 145LEU and 174LEU as geometrically connected residues, whereas these neighbors are missed by the centroid graph.

**Supplementary Table 1**: Neighboring Residues of 147GLY in Rab-33B
Identified by alpha-shape protein graph and centroid graph

| center residue | connected neighbors by alpha-shape protein graph | connected neighbors by centroid graph |
|---|---|---|
| 147GLY | 114TYR,<br>116MET,<br>**145LEU**,<br>146VAL, 148ASN,<br>**174LEU**,<br>175PHE,<br>176GLU, 177THR | **113VAL**,<br>**150CYS**,<br>**163ALA**,<br>114TYR,<br>116MET,<br>146VAL, 148ASN,<br>175PHE,<br>176GLU, 177THR |

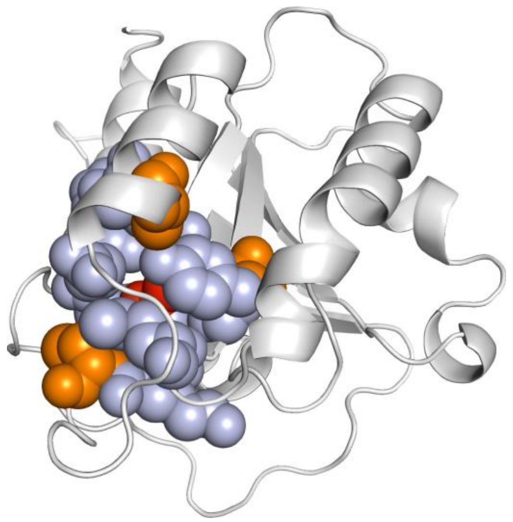

**Supplementary Figure 1**: The center residue 147GLY is labeled in red. Residues connected to 147GLY in the alpha-shape protein graph are labeled in light blue. The centroid graph identifies erroneous neighbors, including 113VAL, 150CYS, and 163ALA, which are labeled in orange. In the visualization, 113VAL, 150CYS, and 163ALA are farther from the center than the neighboring residues (light blue) identified by the alpha-shape graph.

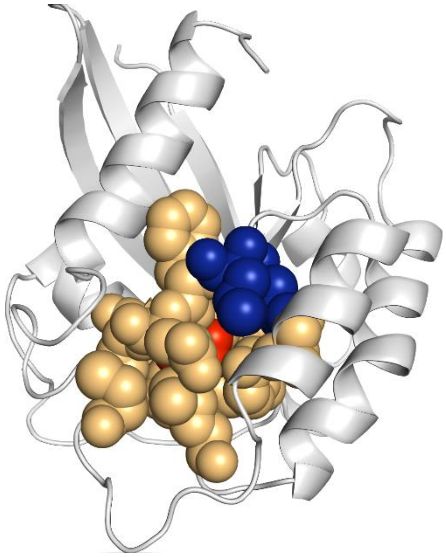

**Supplementary Figure 2**: The center residue 147GLY is labeled in red. Residues connected to 147GLY in the centroid graph are labeled in light orange. The centroid graph fails to capture neighbors 145LEU and 174LEU, which are labeled in blue. In the visualization, 145LEU and 174LEU are relatively close to the center residue.

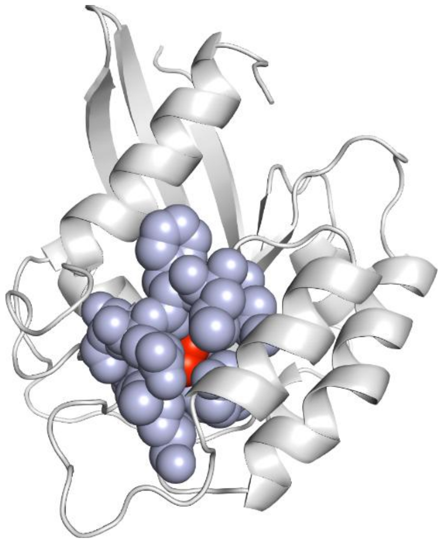

**Supplementary Figure 3**: Connected residues defined by the alpha-shape protein graph. The center residue 147GLY is labeled in red. Residues connected to 147GLY are labeled in light blue.

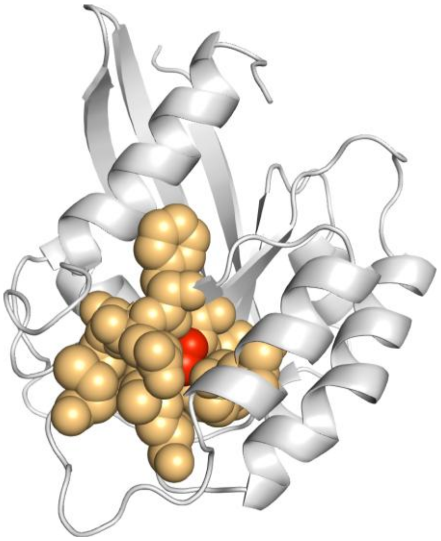

**Supplementary Figure 4**: Connected residues defined by the centroid graph. The center residue 147GLY is labeled in red. Residues connected to 147GLY are labeled in light orange.

##### 1.3 Alpha-shape Protein Graph Reveals Layered Information at Moderate Range

Second, the alpha-shape protein graph reveals precise layered information for neighboring residues at moderate distances, which enables more accurate feature aggregation via *k*-hop message passing in graph neural networks. By contrast, conventional approach typically uses *k*-NN clustering to select the *k* nearest residues (typically 20 or 30 in practice) for 1-hop message passing [2, 4, 9, 10], which ignores intrinsic relative distances to the center residue and does not support explicit *k*-hop message passing.

We continue to use Rab-33B as an example, with 183ASN as the center residue. We selected the 20 closest residues as connected neighbors based on the beta-carbon centroid graph (Supplementary Figure 5). In contrast, the alpha-shape graph precisely reveals that these 20 residues form a three-layer organization relative to the center: 5 residues in the first layer, 13 in the second layer, and 2 in the third layer (Supplementary Figure 6).

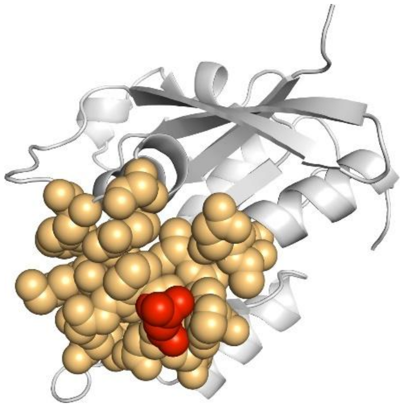

**Supplementary Figure 5**: The *k*-NN ($k$ = 20) neighbors identified by the centroid graph. The center residue 183ASN is labeled in red, and the 20 neighboring residues are labeled in orange.

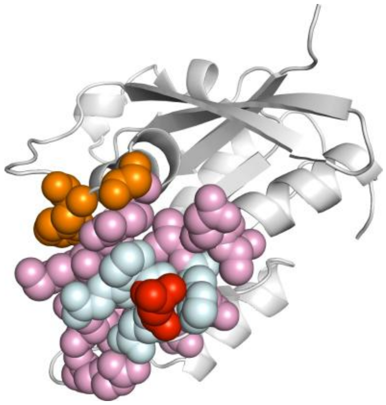

**Supplementary Figure 6**: Layered information for the same $k$ = 20 neighbors revealed by the alpha-shape graph. The center residue 183ASN is labeled in red. Five first-layer residues are labeled in cyan, 13 second-layer residues are labeled in pink, and 2 third-layer residues are labeled in orange.

#### 2. Features

##### 2.1 Structure-derived features for wild-type residue

**bk** denotes the backbone, **sc** denotes the side chain, and **residue** denotes residue-level properties that summarize backbone and side-chain information for the same feature.

**Biophysical properties**: the number of salt bridges for polar residues, backbone dihedral angles, and solvent-accessible surface area (SASA) for the backbone, side chain, and residue. [saltbridge, phi, psi, bk SASA, sc SASA, resi SASA]

**Biochemical properties:** 16 types of atomic interactions between the wild-type residue and its first-layer neighboring residues. [CC, NN, OO, SS, CN, CO, CS, NO, NS, OS, NC, OC, SC, ON, SN, SO] Twenty residue-composition types (contact profiles) in the first, second, and third layers relative to the center residue. [ALA L1, ALA L2, ALA L3, ARG L1, ARG L2, ARG L3, ASN L1, ASN L2, ASN L3, ASP L1, ASP L2, ASP L3, CYS L1, CYS L2, CYS L3, GLN L1, GLN L2, GLN L3, GLU L1, GLU L2, GLU L3, GLY L1, GLY L2, GLY L3, HIS L1, HIS L2, HIS L3, ILE L1, ILE L2, ILE L3, LEU L1, LEU L2, LEU L3, LYS L1, LYS L2, LYS L3, MET L1, MET L2, MET L3, PHE L1, PHE L2, PHE L3, PRO L1, PRO L2, PRO L3, SER L1, SER L2, SER L3, THR L1, THR L2, THR L3, TRP L1, TRP L2, TRP L3, TYR L1, TYR L2, TYR L3, VAL L1, VAL L2, VAL L3]

**Geometric properties** (one-hot encoded): [bk surface, sc surface, bk poc, sc poc, bk buried, sc buried, resi buried, resi poc, resi surface]

##### 2.2 Static Biophysical Changes

Atomic changes and the BLOSUM62 score between the mutant residue and the wild-type residue. [delta backbone C, delta sidechain C, delta sidechain N, delta sidechain O, delta sidechain S, blosum62 score]

##### 2.3 Empirical Capacity Changes

We used the maximum, 99th percentile, and mean to measure the empirical capacity of each of the 20 amino acids, and then computed the differences between the mutant and wild-type residues. These empirical capacities include biochemical and biophysical properties (excluding dihedral angles).

#### 3. Grid Search for Hyperparameters

**Supplementary Table 2. Hyperparameter search in G-SPRI**

| **Hyperparameter** | **Hyperparameter Space** | **Chosen Value** |
|---|---|---|
| Hidden channels (1-hop message passing) | [1024, 512, 384, 256, 192, 128, 64] | 128 |
| Hidden channels (2-hop message passing) | [512, 256, 128, 64, 32, 16] | 512 |
| Hidden channels (3-hop message passing) | [1] (final classification layer) | 1 |
| Rescaling weight for pathogenic class (pos_weight) | [0.2, 0.3, 0.4, 0.5, 0.6, 0.7] | 0.4 |
| Dropout rate | [0.1, 0.2, 0.3, 0.4, 0.5] | 0.5 |
| Weight decay | [0.0005, 0.001, 0.005] | 0.0005 |
| Learning rate | [0.0001, 0.0005, 0.001] | 0.0001 |

#### 4. Detailed performance metrics for binary classification

**Supplementary Table 3. AUPRC for each fold of test dataset**

| | G-SPRI | AlphaMissense | EVE | gMVP | Polyphen-2 |
|---|---|---|---|---|---|
| **Fold 1** | 0.867 | 0.695 | 0.661 | 0.825 | 0.793 |
| **Fold 2** | 0.769 | 0.706 | 0.757 | 0.840 | 0.839 |
| **Fold 3** | 0.877 | 0.703 | 0.766 | 0.829 | 0.813 |
| **Fold 4** | 0.867 | 0.742 | 0.783 | 0.828 | 0.794 |
| **Fold 5** | 0.756 | 0.719 | 0.812 | 0.789 | 0.795 |
| **Average** | **0.827** | **0.713** | **0.756** | **0.822** | **0.807** |
| **std** | **0.059** | **0.018** | **0.057** | **0.019** | **0.020** |

**Supplementary Table 3. AUROC for each fold of test dataset**

| | G-SPRI | AlphaMissense | EVE | gMVP | Polyphen-2 |
|---|---|---|---|---|---|
| **Fold 1** | 0.854 | 0.739 | 0.585 | 0.865 | 0.800 |
| **Fold 2** | 0.755 | 0.751 | 0.624 | 0.847 | 0.828 |
| **Fold 3** | 0.861 | 0.728 | 0.654 | 0.832 | 0.805 |
| **Fold 4** | 0.839 | 0.783 | 0.656 | 0.842 | 0.765 |
| **Fold 5** | 0.713 | 0.775 | 0.709 | 0.827 | 0.788 |
| **Average** | **0.804** | **0.755** | **0.646** | **0.843** | **0.797** |
| **std** | **0.066** | **0.023** | **0.045** | **0.015** | **0.023** |

#### 5. Processing of the TCGA Mutation Profiles

We downloaded the raw TCGA somatic and germline mutation files from the NCI Genomic Data Commons (GDC) database, which used GRCh37 as the reference genome. The somatic mutation files contain pre-annotated information, whereas the germline mutation files are unannotated. To maintain consistency in the transcript and protein isoforms used in this study, we ignored the existing annotations in the somatic files.

We used the standalone VEP tool (version 115) to annotate nucleotide changes and focused on missense mutations [11]. For each gene, we selected a single transcript/protein isoform, obtained a one-to-one mapping between genes and proteins. We used canonical transcript/protein isoforms for 17,438 genes, as defined in Ensembl BioMart. For the remaining 2,547 genes, we selected the first isoform appearing in the VEP output files.

We then used gnomAD v4.1 to remove common germline variants. We set a minor allele frequency (MAF) threshold of 0.001 to define high-frequency alleles. Mutation records were removed if their genomic positions corresponded to any high-MAF alleles in the germline mutation profiles. On average, this process removed 88.9% of germline missense mutations per sample. LiftOver was applied to convert gnomAD chromosome coordinates from GRCh38 to GRCh37.

Additionally, we applied two quality-control steps to improve mutation-call accuracy: (i) Because protein sequences derived from GRCh37-based VEP annotations may be outdated, we used the human reference proteome from UniProt release 2025_03 to correct occasional residue discrepancies. We discarded any mutation record in which the wild-type residue did not match between the Ensembl sequence (from VEP) and UniProt. (ii) We calculated the recurrence of each mutation across the pan-cancer cohort and removed mutations with recurrence greater than 200, because these are likely sequencing or annotation artifacts.

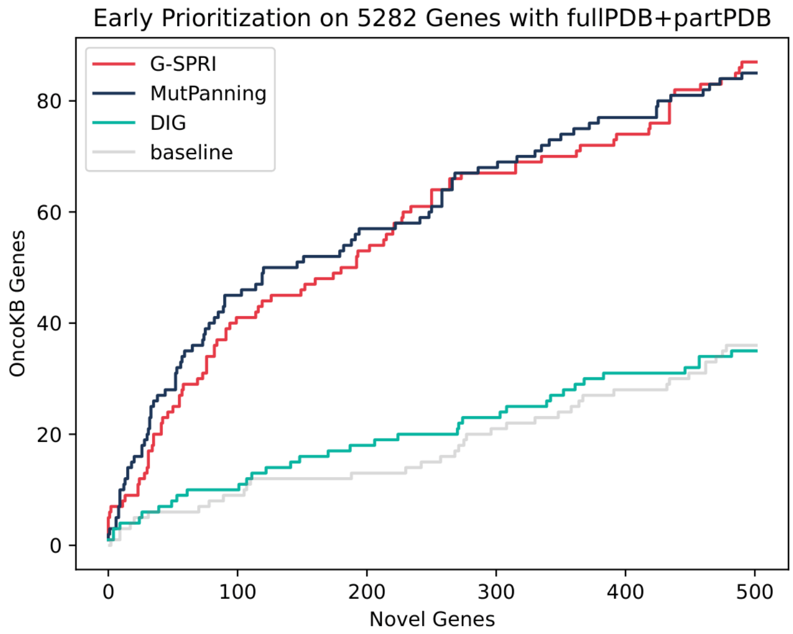


**Supplementary Figure 7**: Early prioritization performance for 5,252 genes with acceptable-quality PDB structures. G-SPRI exhibits performance on par with MutPanning, and shows higher enrichment of OncoKB genes than Dig and baseline.

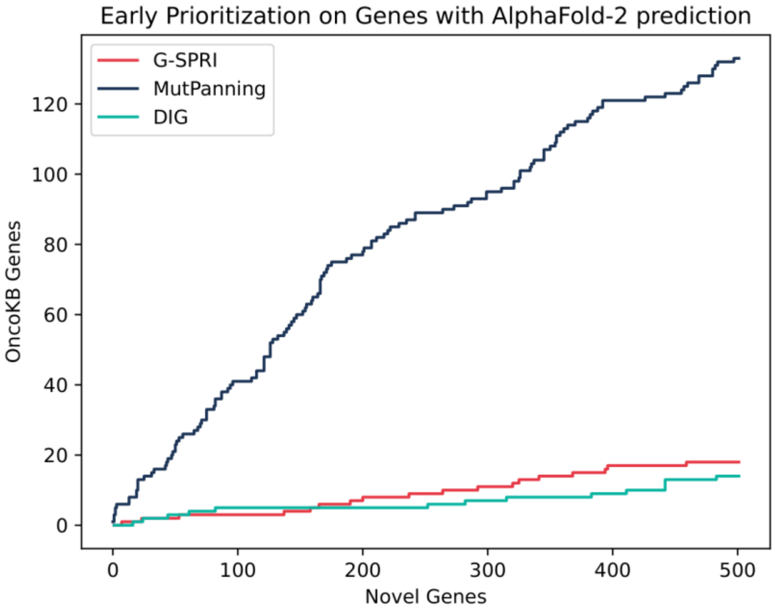


**Supplementary Figure 8**: Early prioritization performance for genes with AlphaFold-2 predicted structures. MutPanning shows the best performance, whereas G-SPRI performs similarly to Dig.
